# Modeling of cytometry data in logarithmic space: when is a bimodal distribution not bimodal?

**DOI:** 10.1101/150201

**Authors:** Amir Erez, Robert Vogel, Andrew Mugler, Andrew Belmonte, Grégoire Altan-Bonnet

**Affiliations:** Immunodynamics Group, Cancer and Inflammation Program, National Cancer Institute, National Institutes of Health, Bethesda, Maryland 20814, USA; IBM T. J. Watson Research Center, Yorktown Heights, New York 10598, USA; Department of Physics and Astronomy, Purdue University, West Lafayette, Indiana 47907, USA; Department of Mathematics, Pennsylvania State University, University Park, Pennsylvania 16802, USA

**Keywords:** bimodal, unimodal, peak, logarithm, gating, FCM, CyTOF

## Abstract

Recent efforts in systems immunology lead researchers to build quantitative models of cell activation and differentiation. One goal is to account for the distributions of proteins from single-cell measurements by flow cytometry or mass cytometry as a readout of biological regulation. In that context, large cell-to-cell variability is often observed in biological quantities. We show here that these readouts, viewed in logarithmic scale may result in two easily-distinguishable modes, while the underlying distribution (in linear scale) is unimodal. We introduce a simple mathematical test to highlight this mismatch. We then dissect the flow of influence of cell-to-cell variability proposing a graphical model which motivates higher-dimensional analysis of the data. Finally we show how acquiring additional biological information can be used to reduce uncertainty introduced by cell-to-cell variability, helping to clarify whether the data is uni- or bimodal. This communication has cautionary implications for manual and automatic gating strategies, as well as clustering and modeling of single-cell measurements.

## Introduction

Flow cytometry (FCM) data typically stretches across several orders of magnitude, with fluorescence intensity *I* readily spanning values between 10^2^ and 10^5^. Although individual cells (events) are recorded as fluorescence intensity *I*, typically what is displayed to the investigator is a binned or smoothed histogram which represents an empirical probability density function (pdf) of the data. Due to the wide span of the data, binning directly the intensity *I* is noisy since there is no single scale one can apply. Instead, when binning FCM data to create a histogram representing a pdf, it is natural to let bin sizes increase as a geometric progression, namely, to evenly bin the logarithm of the fluorescence intensity. As a result, instead of the probability *Q*(*I*) (pdf) of fluorescence intensity *I*, one usually analyzes the probability (pdf) of log *I*, which we denote *P*(log*I*). Indeed, *P*(log*I*) has many advantages: easy display of many orders of magnitude in *I*, easy to model as a two-component log-normal mixture model (as in (1)), and easy to intuitively understand the effect of changing the voltage gain on the flow-cytometer detector photo-multiplier. While such data presentation has been widely adopted in the field of cytometry out of these practical reasons, a rigorous assessment of this log-transformation reveals unwarranted features.

Simply plotting *Q*(*I*) vs. *I* is impractical as most of the data inevitably appears crowded against the *I* = 0 axis. Thus, it is common practice to plot P(log I) or variants thereof which deal with small and negative *I* values introduced by fluorescence compensation (*e.g*. “Logicle” (2), “VLog” (3) and other transformations (4)). Displaying faithfully FCM data is not easy, as the logarithmic scale and fluorescence compensation introduce problems that are easy to miss (5) leading to uncertainty in the number of distinct populations present in the data. Previously, attention has been given to the possibility of effects produced by logarithmic binning (6), contrasting the difference between plotting logarithmic histograms *P*(log *I*) vs. log *I* as opposed to rescaling the x-axis by plotting *Q*(*I*) vs. log I. However, an additional, potentially confusing situation seems to have been overlooked: the possible appearance of a second mode in *P*(log *I*), rendering *P*(log *I*) bimodal, while for the same data only one mode exists in *Q(I*). This is the focus of this work.

When considering biological measurements, *I* is proportional to the *actual* copy number of RNA or proteins. When theoretical considerations are applied to biological systems (such as biochemical dynamics (7, 8, 9, 10, 11, 12), mass-action chemical equilibria, cell-cycle measurements (13) and Hill dose-response curves (14)), it is the copy number itself that is under consideration. Despite that, the logarithm of copy number is an appealing quantity because of its approximately Gaussian statistics, yielding insight into details easily lost if the data were to be analyzed only in linear scale. This leads to a mismatch, where for instance models posed in linear space and data plotted in logarithmic space seem unable to be reconciled without invoking additional effects such as stochastic gene expression noise (7) and cell-to-cell variability (15, 16, 17). Even so, typically one must resort to approximations to analyze noise propagation linearly (8).

The difference between the convenient consideration of the logarithm of abundances and the theoretically-accurate analysis of the linear copy number renders the question of whether *Q*(*I*) has one or two modes (3 extrema) relevant in the following ways: (i) the existence of 1 or 3 extrema is often used to infer the fixed points of a dynamic stochastic biochemical network (7, 9, 1) and other *in silico* methods (18) (ii) extrema are used to define cell-types in automatic (density based) gating and clustering algorithms (19, 20, 21, 22, 23, 24); (iii) The existence of a clearly bimodal distribution is used for manual gating (*e.g*. discerning between activated and un-activated cells) in a way that may appear more robust and compelling than it might truly be. Yet, immunologists have relied on FCM for over forty years to identify new cell populations, often based on manual gating of FCM data. In particular, in cell population enrichment experiments, often such markers are chosen that the (positive) signal is sufficiently well separated from the (negative) background for the two (logarithmic, linear) representations to have no mismatch in their number of peaks. At some level, these experiments succeed in gating the FCM data because typically only these high signal-to-noise markers are used, based on prior knowledge. However, not all FCM experiments enjoy such favorable conditions. Moreover, as the field advances, and more and more markers are acquired, some of these will inevitably have lower signal to noise resolution. The purpose of this manuscript is not to claim that *all* logarithmic gates are problematic. Instead, we caution here of a situation where peaks that appear sufficiently separated in logarithmic space may yet mislead. We propose a simple test to warn if for a given sample, this situations arises.

The rest of this paper is composed of two parts. In the first part, we point out and analyze the situation where a mismatch between the two representations can happen: we formally state the problem using theoretical modeling of FCM data as a mixture of two log-normal distributions (colloquially the ‘‘negative” and ‘‘positive” modes), explicitly show situations where two modes appear in *P*(log *I*) while only one mode exists in *Q*(*I*), and demonstrate this confounding effect on experimental data. In the second part, we analyze the role of cell-to-cell variability in experimental data and show how by measuring a suitable extra dimension one can factor out some of this variability, thus reducing the broadness of the modes sufficiently so as to reduce the mismatch between the two representations. Thus we provide a prescription to design experiments and analyze them so as to resolve the unimodal *vs*. bimodal discrepancy.

## Methods

In this *Methods* section we present, in two detailed sections, the theoretical and experiment techniques used in our study.

### Theory

Given the pdf *P*(log *I*), one can formally derive *Q*(*I*) as (25),

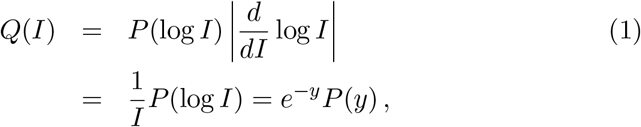

with log *I* ≡ *y*. This relation serves us both in this theoretical section and in the experimental *Results* section, where we estimate the empirical distribution for *Q*(*I*) using this relation. We now address the question: when is *Q*(*I*) unimodal while *P*(log *I*) is multimodal ?

Cytometry data is often amenable to modeling as a log-normal mixture (e.g., (1)). To demonstrate the log/linear mismatch we consider a mixture of two populations, characterized by the distribution of intensity. We define *P*(log *I*) as follows:

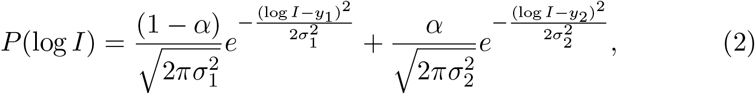

with *y*_1,2_ = log *I*_1,2_ which are the loci of the centers of the left and right Gaussians in log-space, respectively, and *σ*_1,2_ the log-space standard deviations. We then define 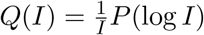 as in Eq. 1.

In Fig. 1, to illustrate with typical measurement values, we set *I*_1_ = 100 and *I*_2_ = 1000 (arbitrary units) and *α* = 0.5, while varying *σ*_1_ = *σ*_2_. This figure presents the three cases we wish to contrast: on the left column, both *P*(log *I*) and *Q*(*I*) are bimodal; in the central column, *P*(log *I*) is bimodal whereas *Q*(*I*) is unimodal; on the right column, both *P*(log *I*) and *Q*(*I*) are unimodal, a situation which we examine in more detail in Eq. 4. Moreover, we see an example where even when both are bimodal (left column), the loci of the modes are different.

**Fig 1.**
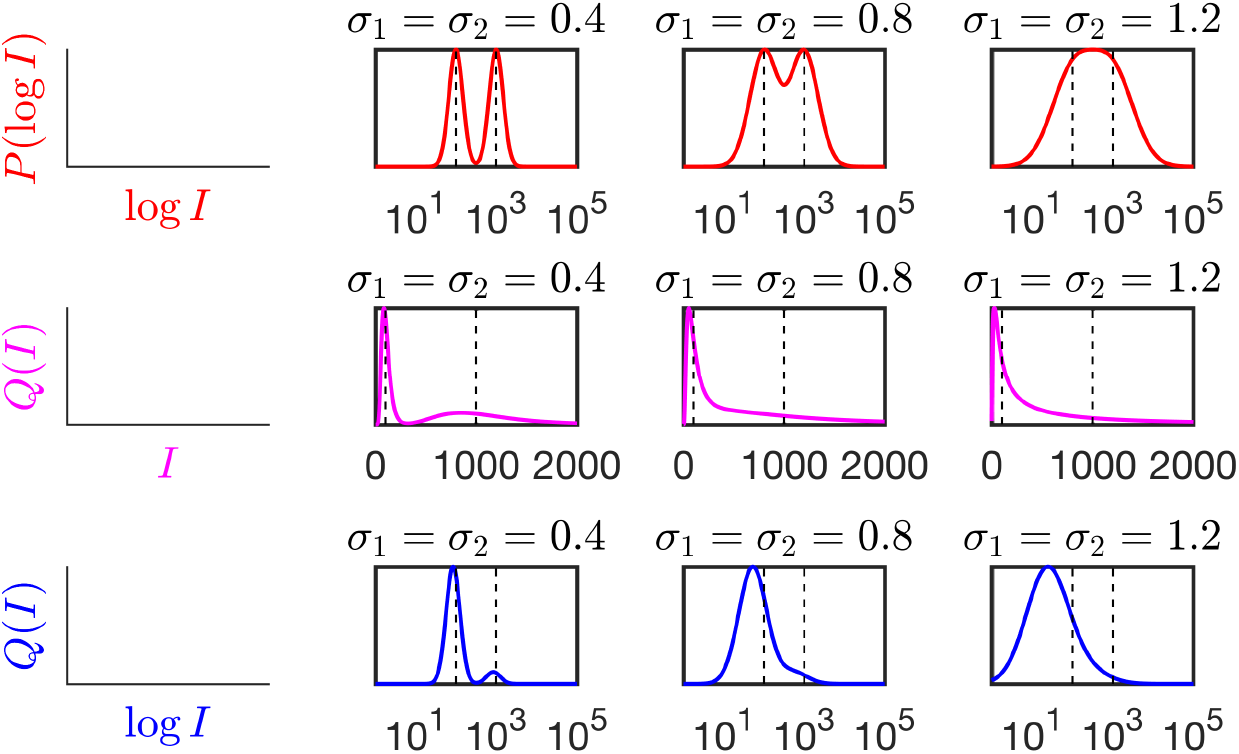
Log-normal mixture showing the mismatch in the number of peaks. Top row, red. *P*(log *I*) vs. log *I* normalized to maximum probability. **Middle row, magenta:** *Q*(*I*) vs. *I* normalized to maximum probability, and plotted on a narrower range. **Bottom row, blue:** *Q*(*I*) vs. log *I* normalized to maximum probability, rescaling the x-axis as in Ref. (6). In the central column where *σ*_1_ = *σ*_2_ = 0.8, P(logI) shows explicit bimodality whereas *Q*(*I*) is unimodal. **Right column:** in this case, the variance *σ*^2^ is large enough (see Eq. 4) that *P*(log *I*) has only one mode even though it is modeled as a mixture. In all cases: *I*_1_ = 100, *I*_2_ = 1000, *α* = 0.5 varying *σ*ı,_2_ = {0.4, 0.8,1.2} (left to right columns, respectively). Leftmost column shows axes.

Determining whether empirical data is multimodal is a difficult task (eg., (26)). Recently, this issue has been revisited in detail (27). In the context of modeling FCM data (18), it was addressed by using Hartigan’s dip test for unimodality (28). Later, as a benchmark to our method we report the results of Hartigan’s test on the experimental data. More details on Harigan’s test can be found in the *Supporting information*.

We define *y* = log *I* and proceed to find the number of extrema for *P*(*y*) and *Q*(*I*). We do this by simple differentiation, demanding that the derivatives of *P* and *Q* equal zero as a definition of extrema. This is achieved by recasting Eq. 2 and its derivatives into the functions *A*(*y*), *B*_1_(*y*), *B*_3_(*y*), defined as,

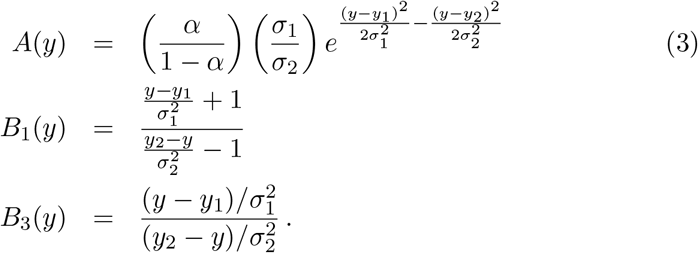

The outcome of this approach is that, to find the extrema of *Q*(*I*), one must find the solutions *y*_\**q*_ for *B*_1_(*y*_\**q*_) = *A*(*y*_\**q*_). Similarly, for the extrema of *P*(log *I*) one must solve *B_3_(y*\*_p_) = *A(y*_p_)*. Full mathematical details on how these are derived can be found in the *Supporting information*.

*A*(*y*) is the ratio of the two Gaussians in Eq. 2 and is therefore always non-negative; this implies that any extremum y* must satisfy *B*_1_(*y*_\**q*_) ≥ 0 and *B*_3_(*y*_\**p*_) ≥ 0. In the *Supporting information*, we derive that the condition for extrema in *Q*(*I*) requires that 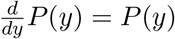 whereas, of course, extremizing *P*(log *I*) sets its derivative to zero, demonstrating the fact that the loci of the modes for *P*(log *I*) and *Q*(*I*) are manifestly different. The region where the log-space distribution shows a second mode occurs when for *B*_3_(*y*_\**p*_) = *A*(*y*_\**p*_) admits three solutions whereas *B*_1_(*y*_\**q*_) = *A*(*y*_\**q*_) admits only one solution. Given that these equations are transcendental, a graphical way to asses the number of solutions is to plot *A*(*y*), *B*_1_(*y*), *B*_3_(*y*) and count the number of times *B*_1_ and *B*_3_ intersect *A*.

In Fig. 2, we present an example of this graphical method. The mismatch between the number of extrema of *P*(log *I*) and *Q*(*I*) is apparent whenever (red curve) *B*_3_(*y*) intersects *A* at 3 points, whereas (blue curve) *B*_1_ (*y*) only intersects *A* once.

**Fig 2.**
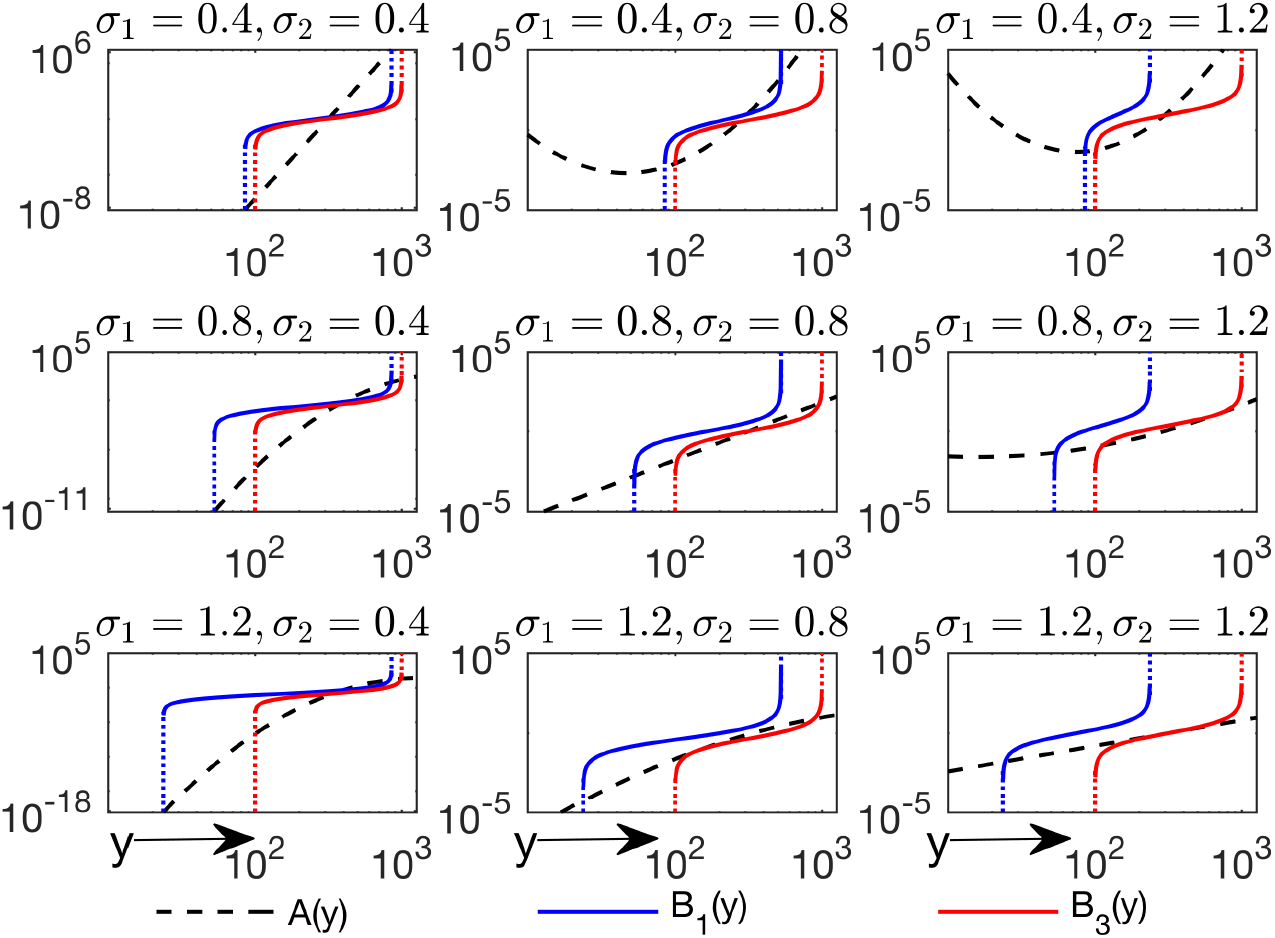
Graphical solution to count the number of extrema. We test our peak counting method, based on the same equations as in Fig. 1, but with additional settings. (Vertical axis represents the functions *A*, *B*_1_, *B*_3_ according to the legend). When the red (blue) curves intersect the dashed black line, *P*(log *I*) (*Q*(*I*)) are extremized. Dashed black: *A*(*y*), blue: *B*_1_(*y*) and red: *B*_3_(*y*). The mismatch between the number of extrema of *P*(log *I*) and *Q*(*I*) is apparent when the red curve intersects *A* at 3 points, whereas the blue curve only intersects *A* once. In both cases, the loci of the extrema are different for the two distributions. The plots along the diagonal (which correspond to the cases in Fig. 1) show the case *σ*_1_ = *σ*_2_ which simplifies *A*(*y*) to a straight line (the general case being a parabola) in these axes.

In the plots along the diagonal, we have *σ*_1_ = *σ*_2_ (as in Fig. 1) which simplifies *A*(*y*) since the quadratic (Gaussian) terms cancel, leaving only an exponential. This leads to a simple criterion to determine whether *P*(log *I*) itself admits one or two modes - previously in Fig. 1(right) we saw an example where *P*(log *I*) is unimodal despite being generated from a mixture. Graphically, we see that for *B*_3_ = *A* to have 3 solutions, log *B*_3_(*y*) has to have a slope less than log *A*(*y*) about the extremum *y*_\**p*_. In other words, 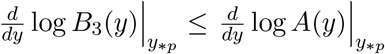, with equality as the threshold between 1 and 3 extrema. This leads to the following intuitive criterion,

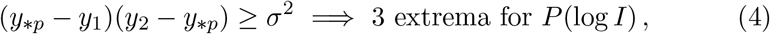

which states that for *P*(log *I*) to appear bimodal, it must have an extremum (*y*_\**p*_) such that the variance of the individual Gaussian components of *P*(log *I*) must be smaller than the distance between *y*_\**p*_ and the Gaussian centers. Substituting for *y*_\**p*_ ≈ log 316, *y*_1_ = log 100 and *y*_2_ = log 1000 and *σ*^2^ = 1.44 we see that the criterion in Eq. 4 is not satisfied and indeed in Fig. 1(right) and Fig. 2(bottom right) we see that *P*(log *I*) has only one mode. A similar condition can be derived for *Q*(*I*), that is, 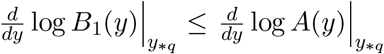, such that,

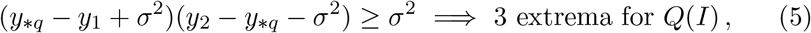

it is, however, hard to compare the two bounds analytically because the *y*_\**p*_ which extremizes *P*(log *I*) is different from the *y*\*_*q*_ which extremizes *Q*(*I*).

### Experimental methods

All data and MATLAB scripts used for the experimental part of this work, together with accompanying MIFlowCyt checklists detailing the data analysis, are freely available online, as follows:

- All MATLAB scripts used to generate the plots and analyze the data in github.com/AmirErez/BimodalLogspaceCytA/tree/master/Scripts
- All data used in the manuscript and accompanying MIFlowCyt checklists come from two experiments:

1. 20140920-OT1-dynamics - used for Fig. 3 in the manuscript.
2. FeinermanScience2008 - previously published in Feinerman et al. Science Vol. 321, Issue 5892, pp. 1081-1084 (2008).

These data with their respective MIFlowCyt checklists are available in github.com/AmirErez/BimodalLogspaceCytA/tree/master/Data

All are accessible in https://github.com/AmirErez/BimodalLogspaceCytA

We briefly sketch the methods for each of the two experiments:

### Experiment 1

Primary mouse T lymphocytes (OT-1) were activated ex vivo and cultured. OT-1 cells were activated by peptide (SIINFEKL) treated and irradiated (3,000 RAD) primary antigen presenting cells (APCs) from a C57BL/6 mouse. OT-1 cells were labeled with an amine-reactive dye, CTV, according to the manufactures protocol (Molecular Probes) for in silico identification. Cells were allowed to rest for one hour after CTV staining, and then distributed in a 96-well v-bottom plate. Cells were then incubated with unique doses of SRC inhibitor Dasatinib at a temperature of 37C for 5 minutes. After which, we added the peptide pulsed APCs (10 APCs to 1 OT-1 T cell) and pelleted mixture of cells by centrifugation for 10 s at 460 rcf at room temperature. The pellet was then incubated at a temperature of 37C for 10 minutes, followed by 15 minutes of chemical fixing with 2% PFA on ice and then permeabilization with ice cold 90% MeOH. Samples were kept at a temperature of -20C until labeling for FCM.

### Experiment 2

C57BL/6N splenocytes were pulsed for 2hr with 100nM SIINFEKL peptide, then irradiated (3000RAD), washed once and used as stimulator/feeder cells. OT-1 lymphocytes were harvested from auxiliary, brachial and inguinal lymph nodes as well as spleen (splenocytes were treated with ACK lysis buffer to remove red blood cells), and mixed with SIINFEKL-pulsed C57BL/6N splenocytes in complete RPMI. After two days, cells were expanded by diluting 2 fold into medium containing 100 pM IL-2. After four days, the cells were again expanded by 2 fold dilution into medium with IL-2. After one more day of culture, cells were harvested and spun through a 1.09 density Ficoll-Paque Plus gradient (GE Healthcare) to remove dead cells. Live cells were recovered, washed twice in complete medium and resuspended at 1 million/ml in complete medium with 100pM IL-2. Cells were used for experiments between 6 and 8 days after primary stimulation. The ppERK response to SRC and MEK inhibition was measured using primary OT-1 T-lymphocytes activated with RMA-S APCs. RMA-S cells were suspended in culture with 1 nM SIINFEKL peptide for 2 h at 37C, 5% CO**2**, and on a rotator to guarantee mixing. During this time we labelled OT-1 cells with an amine-reactive dye, CTV, according to the manufactures protocol (Molecular Probes). We rested the OT-1 cells one hour after CTV staining, and then distributed them in a 96-well v-bottom plate. Each well was given various doses of SRC inhibitor and MEK inhibitor and kept at 37C for 5 min. Following the 5 min exposure to the inhibitors, we added the peptide pulsed RMA-S (10 RMA-S to 1 OT-1 T cell) and pelleted by centrifugation for 10 s at 460 rcf at room temperature. This step guaranteed that both cell types, OT-1 and RMA-S, came into contact. The cells were allowed to activate for 10 min, followed by fixing on ice in 2% PFA, and then permeabilized and stored in 90% MeOH at -20C. APCs were pulsed with serial dilutions of OVA or variant peptides for 2 hr at 37C, then washed with T cell medium at the time of harvest, and resuspended with anti-CD8 (53-6.72)-Fab-coated T cells in their conditioned media in a V-bottom 96-well plate (Corning). Fab-coating was performed 10 min before cell use by incubating T cells with 10 *μg*/ml of Fab fragment. T:APC cell contacts were synchronized using a quick centrifugal spin (10s at 400g). Plates containing T:APC conjugates were placed on a water bath at 37C and incubated for 5 min. Supernatants were then discarded, T:APC conjugates disrupted by vortexing, and cells resuspended in ice-cold 4% paraformaldehyde for 15 min. Cells were then permeabilized with ice-cold 90% methanol for 15 min on ice, and washed twice with FACS buffer.

## Results - experiments

We now apply our analysis to experimental data, by analyzing two different experiments, described briefly in the *Methods* section and in more detail in the MIFlowCyt checklists available online. Both experiments measure the distribution of Extracellular Signal-regulated Kinase (ERK) phosphorylation (ppERK) signaling in CD8+ primary mouse T-cells responding to antigens.

In experiment 1, the T-cells were inhibited by the SRC inhibitor Dasatinib. In Fig. 3(A) we see how a commercially-available analysis software (FlowJo (29)) plots the distribution of ppERK in such an experiment, which clearly shows a bimodal structure. Fig. 3(B) plots those same data, formally 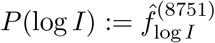, the probability density estimated from the data by taking the logarithm of the intensity of 8751 cells and binning it in bins of size 0.2. Our procedure for estimating *P*(log *I*) gives the two modes as in FlowJo (red dots) whereas *Q*(*I*) (derived using Eq. 1) has a single mode (blue dots). It also agrees with the results of Hartigan’s unimodality p-values, *p_U_*, explained in more detail in the *Supporting information*. We fit *P*(log *I*) as a Gaussian mixture. This is followed in Fig. 3(C) by the same extrema analysis as in Fig. 2, revealing that indeed *Q*(*I*) has a single maximum.

**Fig 3.**
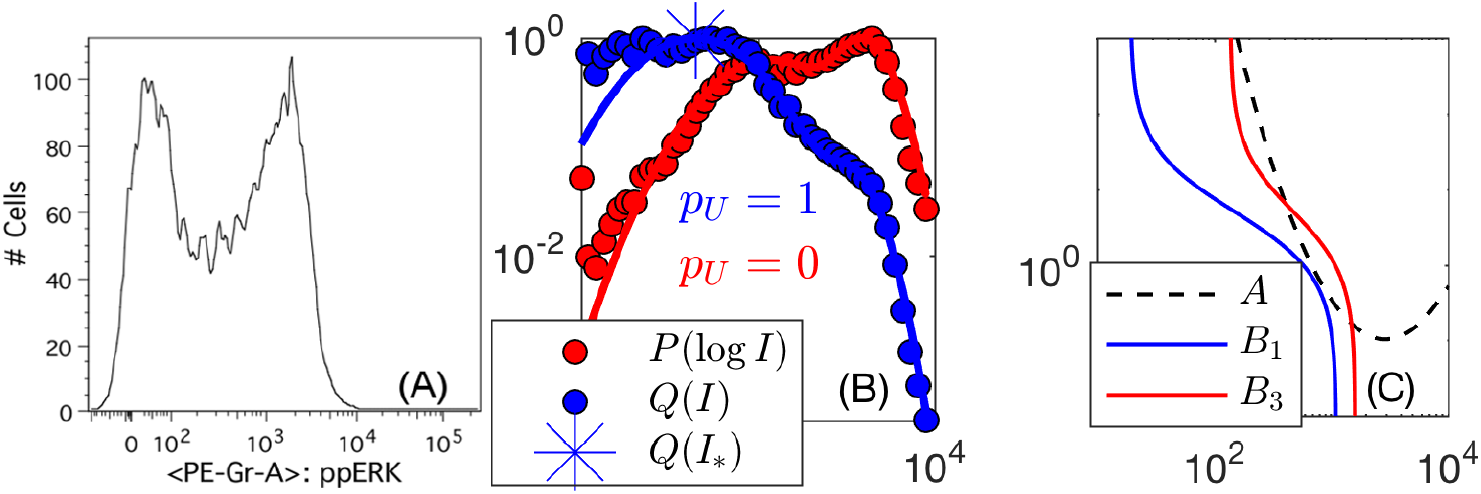
Analysis of experimental data reveals the effect we describe in a real scenario. **(A)** Histogram of ppERK as plotted by FlowJo (29); **(B)** Histograms for P(logI) (red,dots) and *Q*(*I*) (blue,dots) vs. log *I* as estimated from the data binned logarithmically. Note that plotting *Q*(*I*) vs. log *I* is somewhat unusual but allows both P and Q to be plotted on the same axis. The red line shows the result of fitting P(logI) to a gaussian mixture model (Eq. 2), and the blue line is the estimate for *Q*(*I*) from *P*(log *I*) according to Eq. 1. The blue star indicates the location of the only maximum for *Q*(*I*) obtained from Eq. 8, despite the obvious two maxima in P(logI) (red). Hartigan’s unimodality p-values for log *I* (red) and *I* (blue) are taken directly from the data without binning, corroborating that log *I* is bimodal whereas *I* is unimodal. **(C)** (Vertical axis represents the functions *A*, *B*_1_, *B*_3_ according to the legend). Graphic solution of the extrema conditions as in Fig. 2 explicitly reveals the three solutions for *P*(log *I*_*_) (red line intersects black dashed) as opposed to the single solution for *Q*(*I*_*_) (the blue line intersects the black dashed line up beyond the plotted area, solution also plotted as blue star in the middle plot), indicating that *Q*(*I*) has only one mode.

Given the prevalence and success of manual gating of FCM data in many situations, we wondered whether gating in logarithmic scale could be justified *a posteriori*, based on biological knowledge. In the final part of our analysis, we aimed to include additional information such that the biological significance of the distributions in our single-cell measurements is better captured.

We proceed to analyze experiment 2, these data were taken from Ref. (15). We briefly sketch what follows: by considering both ERK1 and ppERK levels, (the two being biologically related and experimentally correlated), we demonstrate that dividing one by the other ameliorates the mismatch between the logarithmic and linear representations. This is done by examining their *joint* probability density function *P*_2_; we suggest a probabilistic graphical model that describes the correlations in the system and their dependence on cell-to-cell variability; we suggest a way to normalize for cell-to-cell variability, such that the mismatch (resulting from increased uncertainty) between logarithmic and linear gates is resolved. We then generalize our example and provide a simple experimental and theoretical framework that allows less ambiguous determination of positive/negative gates.

In Fig. 4(A), we show the experimental data we will use in our proposed solution. We show a heat map of the *joint* distribution of ppERK (*I_ppERK_*) and total ERK1 (*I*_*ERK*1_) expression in mouse CD8+ T-cells. We estimate their joint distribution by using a kernel density estimator (30), which is used only for presentation purposes and for demonstrating their conditional independence (Fig. 6). Notably, whereas the two modes of ppERK significantly overlap when plotted in the marginal distribution *P*(log *I_ppERK_*), ppERK expression correlates with total ERK1 levels in their joint distribution *P*_2_(log *I_ppERK_*, log *I*_*ERK*1_). The relation in Eq. 1 can be generalized to *P*_2_(log *I_ppERK_*, log *I*_*ERK*1_) (see, eg., (25)), however, it will not be used in this manuscript.

**Fig 4.**
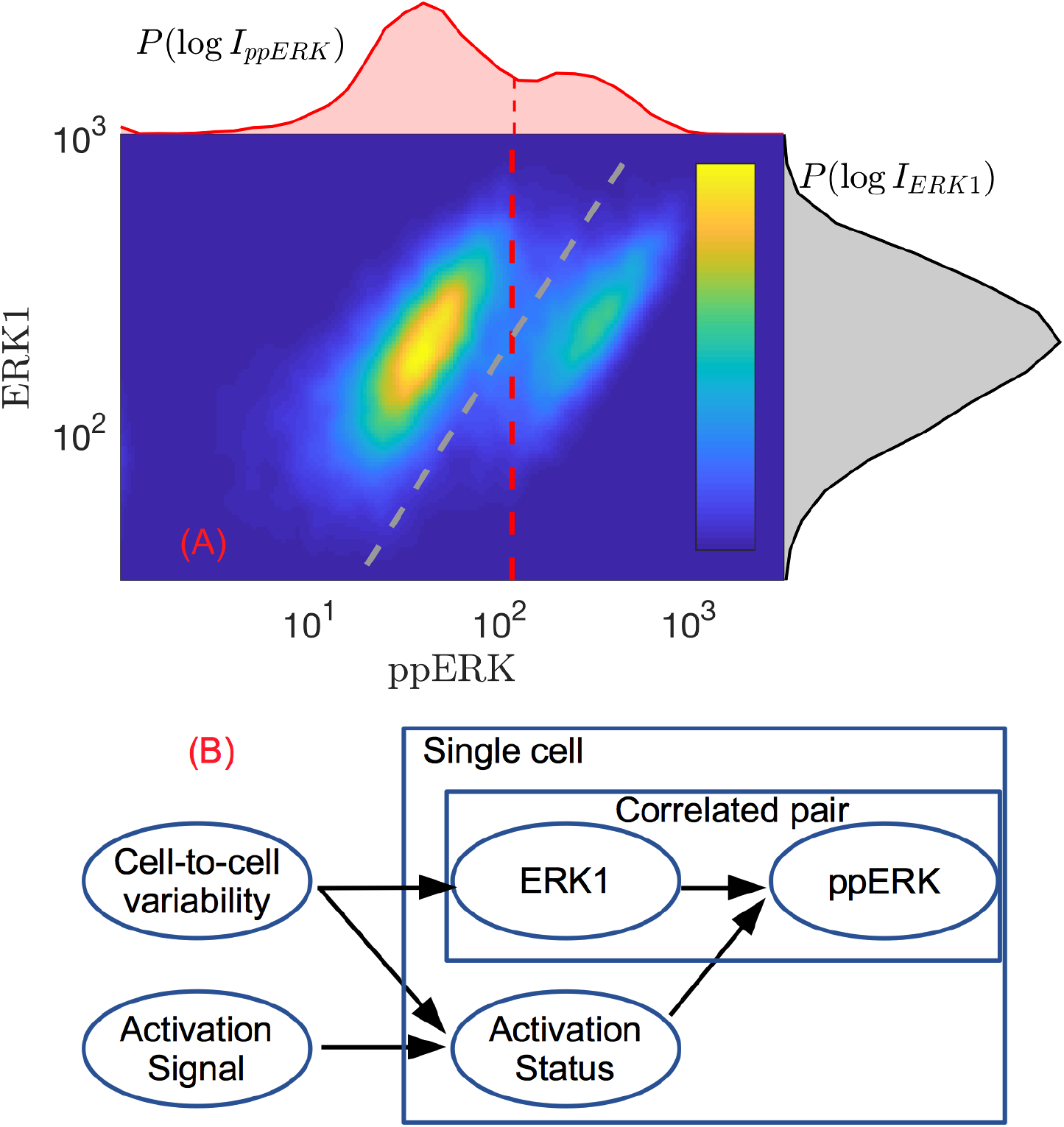
Analysis of experimental data with two correlated measurements. **(A)** The joint distribution *P*_2_(log *I_ppERK_*, log *I*_*ERK*1_) as a heat map with its marginals plotted on the top and on its right. The correlation between ppERK and ERK1 levels is clear in the data. Dashed red (grey) lines are proposed manual gates according to the marginal (joint) distributions *P*(log *IppERK*) (*P***2** (log *IppERK*, log *I*_*ERK*1_)). **(B)** Bayesian network depicted as a graphical model to show the flow of influence on the measurement of ppERK. The pair ERK1 and ppERK are in a template to suggest that there exist other pairs of correlated observables that depend on activation status and cell-to-cell variability.

**Fig 5.**
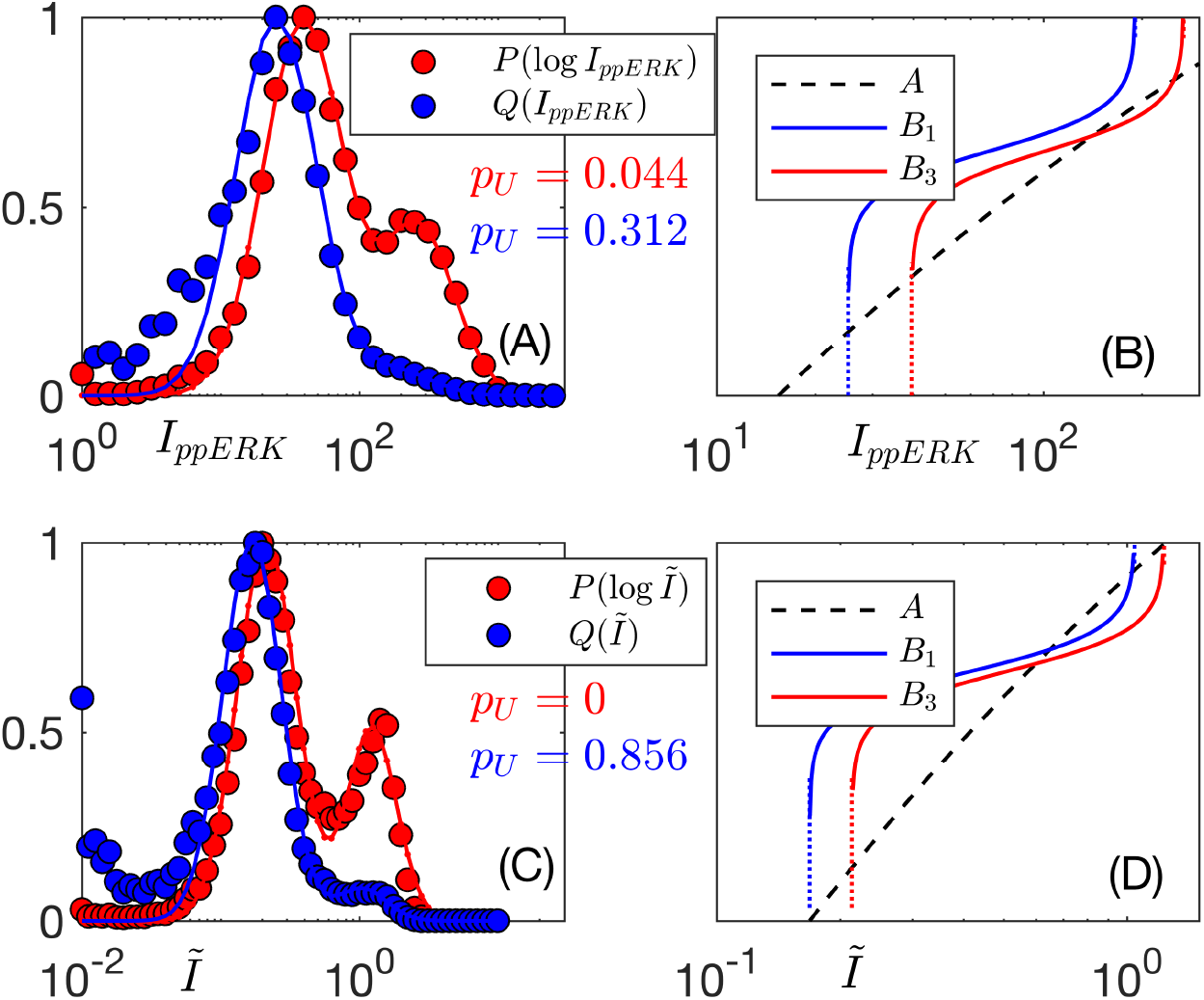
By dividing ppERK readings by ERK1, we can ameliorate the mismatch between the two representations. **(A)** *P*(log *I_ppERK_*) and *Q*(*I_ppERK_*) vs. log *I_ppERK_* (dots: data, lines: Gaussian mixture fit) together with **(B)** their extrema analysis, showing that the second mode in log ppERK does not exist if the data is linearly binned. **(C)** The same treatment but for *Ĩ* = *I_ppERK_*/*I*_*ERK1*_, **(D)** shows that both *I* and log *I* have two modes, thus normalizing ppERK levels by total ERK1 maintains the bimodal structure both in *P*(log *Ĩ*) and in *Q*(*Ĩ*). In (B) and (D), the vertical axis represents the functions *A*, *B*_1_, *B*_3_ according to the legend.

**Fig 6.**
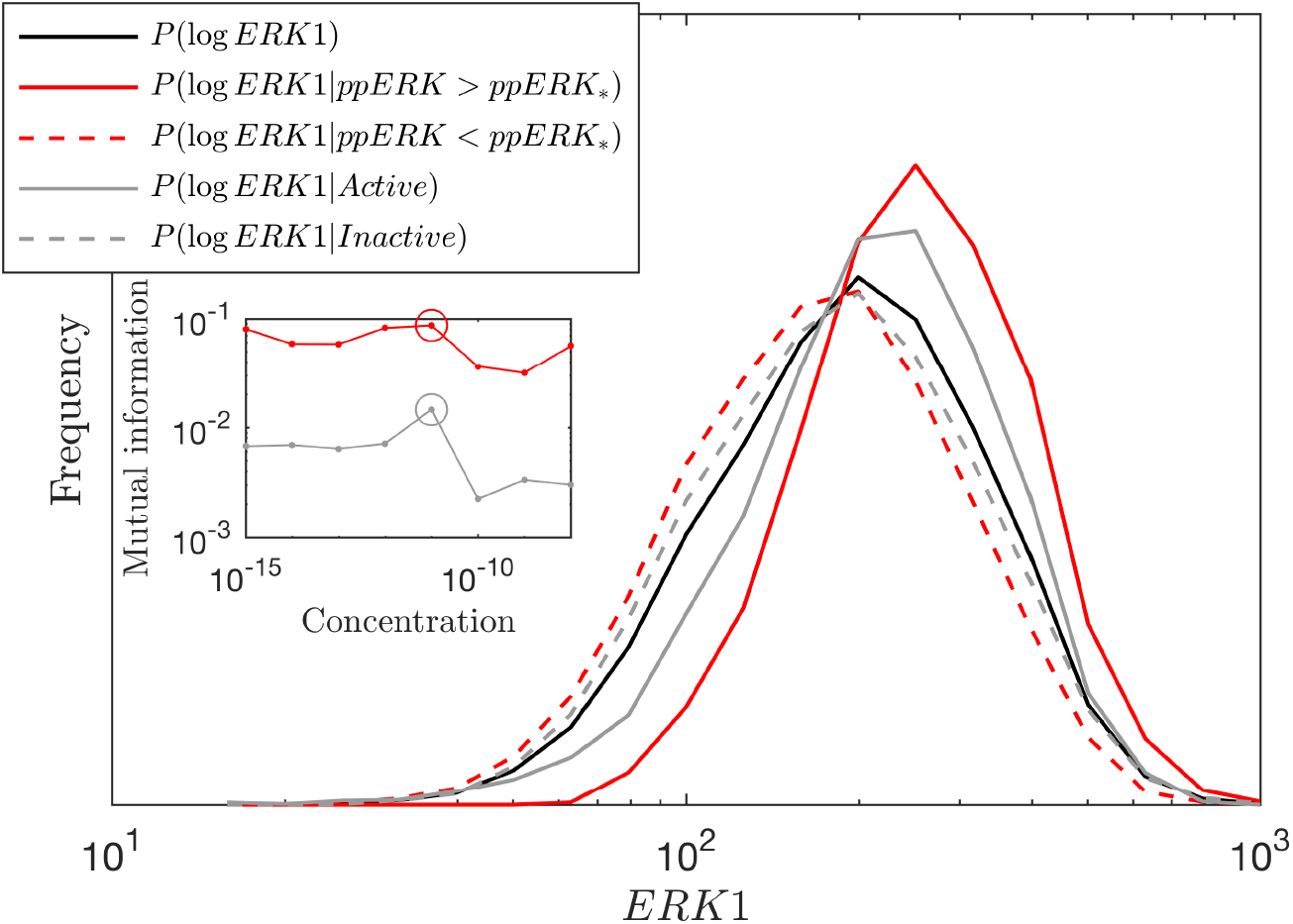
Test for weak dependence of ERK1 and activation status. Whereas independence implies that *P*(log *ERK*1) = *P*(log *ERK*1|*Activation*), in fact there is a weak dependence between them regardless of whether we employ a vertical (red, defined by threshold value *ppERK*_*_) or a diagonal (grey) gate (the diagonal gate showing a weaker dependence); this is observed directly by noting that the different distributions in Fig. 6 do not lie on top of each other. To quantify this difference, the inset shows the mutual information between *P*(log*ERK*1) and *P*(log *ERK*1|*Activation*), with the circle pointing out the particular concentration of stimulus (out of all concentrations used in Ref. (15)) we chose to plot in this example. The chosen concentration has the highest mutual information, *i.e*., the lowest ability to discern between the active and inactive states.

To estimate the marginal densities *P*(log *I_ppERK_*) and *P*(log *I_ERK1_*) we bin the log-intensity of 24449 cells. Each cell’s state encodes another *latent* variable, its activation status - which tells if the cell has been successfully activated by the stimulus. To deduce the activation status, it is common practice to use manual gating of the data by drawing a boundary between the active and inactive states. To account for the correlation between ERK1 and ppERK we consider two manual gating strategies: (i) perpendicular gating (dashed red) according to *P*(log *I_ppERK_*) with *I_ppERK_* > *ppERK*_*_ considered an activated cell, and (ii) diagonal gating according to the apparent correlation in *P*_2_ (dashed grey). We set the diagonal gate with a slope of unity, meaning that we take the dividing line, reflecting proportionality *ppERK* ∝ *ERK1*, as a good way to partition the two states. We define ‘‘Inactive” to the left of the dashed line, and ‘‘Active” to the right of it.

To understand the structure of these data, it is important to characterize explicitly the dependency structure of our observables (ERK1, ppERK), the latent activation status, and the influence of external factors on these three. The existence of two peaks in ppERK which appear distinct from each other but correlated with ERK1 levels, guides us to use a Bayesian network to capture these features in the data as a graphical model. First - we test whether ERK1 and the cell’s activation status are independent. In Fig. 6 we see that whereas independence implies that *P*(log *ERK*1) = *P*(log *ERK*1|*Activation*), in fact there is a weak dependence between them regardless of whether we employ a vertical (red) or a diagonal (grey) gate (the diagonal gate showing a weaker dependence). The weak dependence between activation state and ERK1 levels is reasonable, if we account for cell-to-cell variability, since for a given stimulus some cells inevitably respond differently from the typical cell (31).

We summarize the causal structure for this system in Fig. 4(B) which depicts a probabilistic graphical model (32) of the flow of influence from cell-to-cell variability and activation signal, to ERK1 levels and activation status, and finally to the distribution of ppERK. We depict the pair ERK1-ppERK in a template (rectangles), to suggest to the reader the existence of multiple other pairs. Thus, by formalizing the flow of influence of cell-to-cell variability on the observed fluorescence signal we provide reasoning for what follows.

We proceed to show how to better resolve the log-space peak; this recipe, together with the model in Fig. 4(B) can be used *a priori* in automatic gating and clustering algorithms to prevent some of the mismatch between logarithmic and linear binning strategies. For stochastic modeling, such a structure presents an opportunity to analyze the structure and propagation of noise in the system (8, 33).

We treat the broadness of ppERK modes as generated by cell-to-cell variability in total ERK1 content - a reasonable assumption since the noise in the phosphorylation of ERK is negligible in comparison (34). We further neglect the indirect influence between activation status and ERK1 levels due to its weakness (checked in Fig. 6). Thus we approximate that the conditional independence between ERK1 and activation status (given that both are influenced by cell-to-cell variability) is true independence. This implies an approximately linear relation *I_ppERK_* ∝ *I*_*ERK*1_ given activation status. We define the normalized intensity *Ĩ* = *I*_*ppERK*_/*I*_*ERK*1_ as the ratio of ppERK to ERK1 intensity, thereby eliminating the linear dependence of ppERK on ERK1 levels and reducing uncertainty due to cell-to-cell variability. The resulting *P*(log *Ĩ*) may boast a sufficiently reduced noise in ppERK such that a clear bimodal signature appears regardless of logarithmic or linear binning of *Ĩ*. In Fig. 5 we show such an example, where in Fig. 5(A,B), *P*(log *I*) and *Q*(*I*) do not agree on the number of modes, whereas in Fig. 5(C,D) *Ĩ* = *I_ppERK_*/*I*_*ERK*1_ do agree. These data have order 25,000 events and so, similarly to Fig. 7, Hartigan’s test may not identify the number of peaks correctly, as is indicated in the *p_u_* values.

Thus we demonstrate how by suitably accounting for cell-to-cell variability one can reduce the measured noise so as to circumvent the mismatch in the number of modes between the logarithmic and linear treatment. For testing bimodality, whereas our method relies on fitting a Gaussian mixture, existing statistical tests, eg., Hartigan’s test, require no fitting yet may lack statistical power when applied to typical experimental situations.

## Discussion

We conclude our analysis by making a more intuitive argument. Above, we have demonstrated the theoretical and experimental existence of a situation where *P*(log *I*) has two modes when *Q*(*I*) has only one. But how does this *come about* ? Intuitively, by binning in logarithmic scale, we are effectively making the bin sizes grow as *I* increases. A larger bin can only lead to a higher count in that bin and so we might stumble upon a regime where this creates a quasi-mode. This situation and the same logic applies also when using more advanced logarithmic-like transformations, (eg., Logicle (2) and Vlog (3)), as is demonstrated in Fig. 3A.

The scale-dependent bimodality as demonstrated in Fig. 3 and Fig. 5(A,B) may be not uncommon. Specifically, one must take extra care when attempting to manually gate, automatically cluster or build dynamical models which rely on an apparent bimodal structure, as it might depend on whether the data was log-transformed or not. This becomes increasingly relevant as cytometry moves forward to higher dimensional measurements which become tractable only with automatic gating schemes. Instead, one might consider plotting *Q*(*I*) on the log-log scale, a presentation which preserves the number of maxima, at the expense of the measure of the distribution. It is possible to ameliorate the mismatch between the two scales, as we demonstrate in Fig. 5(C,D), if one can simultaneously measure correlated observables (in our example, ppERK and ERK1). This allows to control for cell-to-cell variability, increasing the resolution of the data. Recently, this favorable scenario has become more attainable with the introduction of mass cytometry - where one can rely on a large number of channels without compromising the FCM panel. Based on the analysis carried out in this paper, we conjecture that such extra channels, chosen wisely, can provide automatic clustering/gating algorithms the right information needed to make more reliable clustering and population defining. This is a simple way to introduce knowledge of the biological structure of the data into otherwise objective clustering algorithms, without compromising their objectivity.

We propose and test some features of a graphical model that captures the structure of such dependencies and the propagation of noise from cell-to-cell variability into the observed fluorescence signal. The graphical model is used to motivate our suggestion of how to ameliorate the mismatch between the linear and logarithmic representations. Moreover, it is potentially useful for those interested in correctly capturing these dependencies for automatic gating and clustering algorithms.

Logarithmic transform of FCM intensity values is the usual practice, but for some observations, eg., the FSC and SSC channels, linear scale is typically used. In such cases, there is no danger of mismatch between presentation. When FCM data are logarithmically distributed, though we caution on the use of *P*(log *I*), we find it remarkable how well the distribution of biological quantities resembles a log-normal mixture.

## Acknowledgments

This work was supported by Human Frontier Science Program grant LT000123/2014 (Amir Erez) and by the Intramural Research Program of the NCI, NIH.

## Conflicts of Interest

The authors declare that there are no conflicts of interest.

## Supporting information

### Finding the extrema of a log-normal mixture

It is possible to discern between one or three extrema of the distribution *P*(log *I*), corresponding to one or two modes (respectively) by counting the number of solutions for 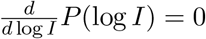. Similarly, 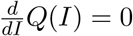 can have either one or three solutions. This raises the possibility of there being three extrema (two modes) for *P*(log*I*) whereas only one mode in *Q*(*I*). We explicitly evaluate the extrema of the mixture of log-normal distributions by solving,

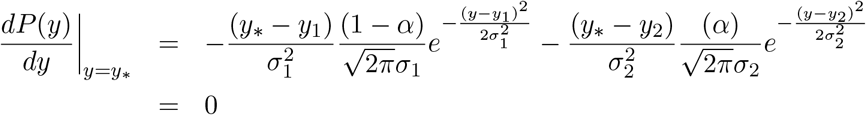

which by algebraic rearrangement,

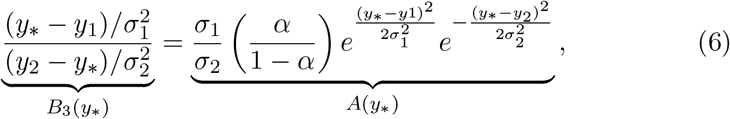

provides a more transparent form.

Here we refer to the left-hand-side and right-hand-side as *B*_3_(*y*) and *A*(*y*), respectively. In like, computing the extrema of the linear scale distribution amounts to 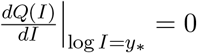, which by change of variable is equivalent to,

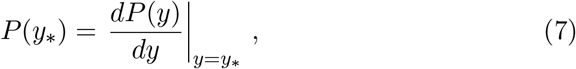

and by substitution,

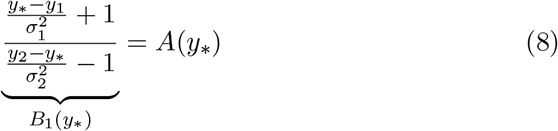

We refer to the quantity on the left-hand-side, *B*_1_(*y*).

### Hartigan’s dip test

Briefly, Hartigan’s dip statistic measures the maximum difference between the empirical distribution and the unimodal distribution that minimizes that maximum difference. This is compared to the appropriate null distribution which is, in this case, the uniform distribution, to give *p_u_*, a p-value for unimodality.

As a check for the predictive power of Hartigan’s test with regards to experimental data, we apply it on a log-normal mixture comparing its predictive power as a function of the number of tests and number of events in each test (35). For this, we draw events from *Q*(*I*) by using MATLAB’s *lognrnd* function for each of the two log-normal distributions. In Fig. 7, we test it on the situation in Fig. 2(top,middle) in which *Q*(*I*) is weakly bimodal, meaning that its bimodality is nearly marginal (*σ***1** = 0.4 and *σ*_2_ = 0.8). The Hartigan probability of unimodality (*p_u_*) is not sensitive to the number of bootstrap tests in a reasonable range but becomes strongly predictive of the (weak) bimodality only when there are more than 10^5^ events. Such an abundance of cells may not always be available in typical FCM data, especially for sub-populations which have been selected (gated) and may comprise only a small fraction of all the cells acquired. Fig. 7 supports that in such weakly bimodal situations, Hartigan’s p-value should be treated cautiously, a situation which we encounter in Fig. 5.

**Fig 7.**
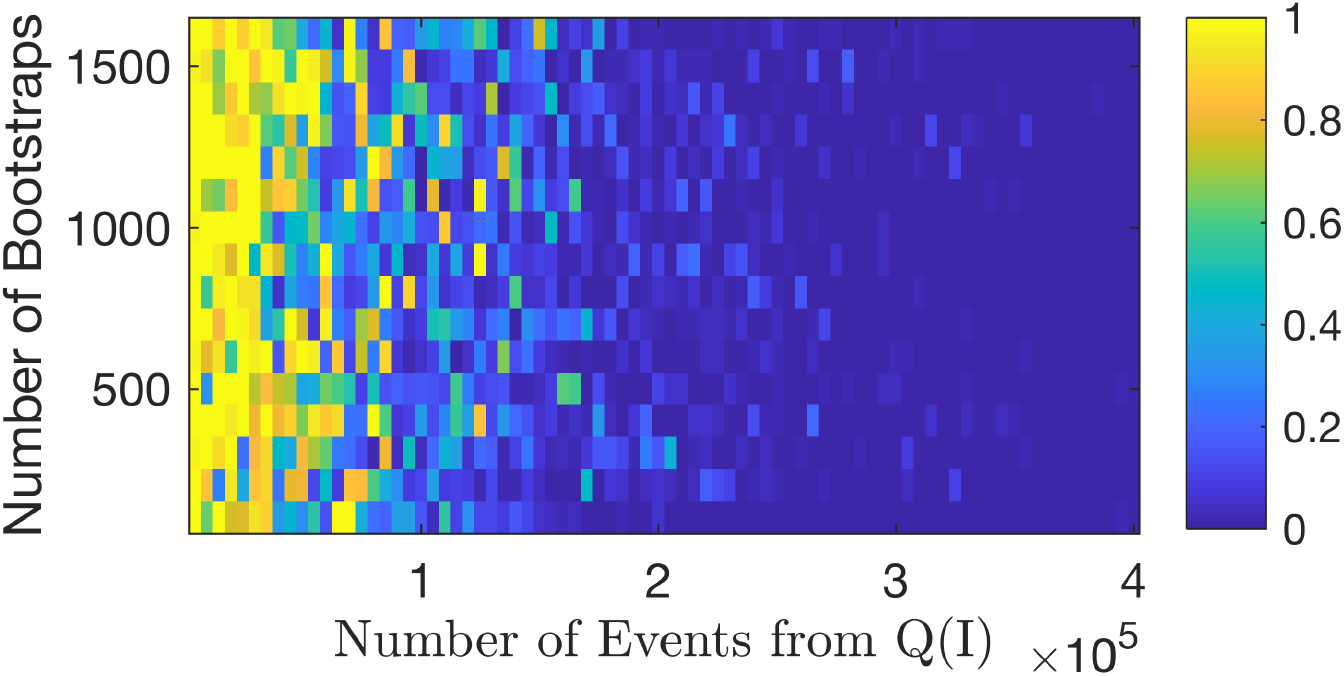
Testing Hartigan’s p-value *p_u_* for unimodality on a weakly bimodal log-normal mixture. The heat-map shows *p_u_* as a function of the number of bootstrap tests and number of events in each test, for *σ*_1_ = 0.4 and *σ*_2_ = 0.8 as in Fig. 2(top,middle). With less than ~ 10^5^ events, Hartigan’s test for this case may misidentify the number of peaks.

